# Creating Artificial Human Genomes Using Generative Models

**DOI:** 10.1101/769091

**Authors:** Burak Yelmen, Aurélien Decelle, Linda Ongaro, Davide Marnetto, Corentin Tallec, Francesco Montinaro, Cyril Furtlehner, Luca Pagani, Flora Jay

## Abstract

Generative models have shown breakthroughs in a wide spectrum of domains due to recent advancements in machine learning algorithms and increased computational power. Despite these impressive achievements, the ability of generative models to create realistic synthetic data is still under-exploited in genetics and absent from population genetics.

Yet a known limitation of this field is the reduced access to many genetic databases due to concerns about violations of individual privacy, although they would provide a rich resource for data mining and integration towards advancing genetic studies. In this study, we demonstrated that deep generative adversarial networks (GANs) and restricted Boltzmann machines (RBMs) can be trained to learn the high dimensional distributions of real genomic datasets and create high quality artificial genomes (AGs) with none to little privacy loss. To illustrate the promising outcomes of our method, we showed that (i) imputation quality for low frequency alleles can be improved by augmenting reference panels with AGs, (ii) scores obtained from selection tests on AGs and real genomes are highly correlated and (iii) AGs can inherit genotype-phenotype associations. AGs have the potential to become valuable assets in genetic studies by providing high quality anonymous substitutes for private databases.

## Introduction

Availability of genetic data has increased tremendously due to advances in sequencing technologies and reduced costs (Mardis 2017). The vast amount of human genetic data is used in a wide range of fields, from medicine to evolution. Despite the advances, cost is still a limiting factor and more data is always welcomed, especially in population genetics and genome-wide association studies (GWAS) which usually require substantial amounts of samples. Partially related to the costs but also to the research bias toward studying populations of European ancestry, many autochthonous populations are under-represented in genetic databases, diminishing the extent of the resolution in many studies (Cann 2002; Popejoy and Fullerton 2016; Mallick et al. 2016; Sirugo et al. 2019). Additionally, a huge portion of the data held by government institutions and private companies is considered sensitive and not easily accessible due to privacy issues, exhibiting yet another barrier for scientific work. A class of machine learning methods called generative models might provide a suitable solution to these problems.

Generative models are used in unsupervised machine learning to discover intrinsic properties of data and produce new data points based on those. In the last decade, generative models have been studied and applied in many domains of machine learning (Libbrecht and Noble 2015; Zhang et al. 2017; Rolnick and Dyer 2019). There have also been a few applications in the genetics field (Davidsen et al. 2019; Liu et al. 2019; Tubiana et al. 2019; Shimagaki and Weigt 2019), one specific study focusing on generating DNA sequences via deep generative models to capture protein binding properties (Killoran et al. 2017). Among the various generative approaches, we focus on two of them in this study, generative adversarial networks (GANs) and restricted Boltzmann machines (RBMs). GANs are generative neural networks which are capable of learning complex data distributions in a variety of domains (Goodfellow et al. 2014). A GAN consists of two neural networks, a generator and a discriminator, which compete in a zero-sum game (Supplementary Figure 1). During training, the generator produces new instances while the discriminator evaluates their authenticity. The training objective consists in learning the data distribution in a way such that the new instances created by the generator cannot be distinguished from true data by the discriminator. Since their first introduction, there have been several successful applications of GANs, ranging from generating high quality realistic imagery to gap filling in texts (Ledig et al. 2017; Fedus et al. 2018). GANs are currently the state-of-the-art models for generating realistic images (Brock et al. 2018).

A restricted Boltzmann machine, initially called Harmonium is another generative model which is a type of neural network capable of learning probability distributions through input data (Smolensky 1986; Teh and Hinton 2001). RBMs are two layer neural networks consisting of an input (visible) layer and a hidden layer (Supplementary Figure 2). The learning procedure for the RBM consists in maximizing the likelihood function over the visible variables of the model. This procedure is done by adjusting the weights such that the correlations between the visible and hidden variables on both the dataset and sampled configurations from the RBM converge. Then RBM models recreate data in an unsupervised manner through many forward and backward passes between these two layers (Gibbs sampling), corresponding to sampling from the learned distribution. The output of the hidden layer goes through an activation function, which in return becomes the input for the hidden layer. Although mostly overshadowed by recently introduced approaches such as GANs or Variational Autoencoders (Kingma and Welling 2013), RBMs have been used effectively for different tasks (such as collaborative filtering for recommender systems, image or document classification) and are the main components of deep belief networks (Hinton and Salakhutdinov 2006; Hinton 2007; Larochelle and Bengio 2008).

Here we propose and compare a prototype GAN model along with an RBM model to create Artificial Genomes (AGs) which can mimic real genomes and capture population structure along with other characteristics of real genomes. We envision two main applications of our generative methods: (i) improving the performance of genomic tasks such as imputation, ancestry reconstruction, GWAS studies, by augmenting genomic panels with AGs serving as proxies for private datasets, (ii) demonstrating that a proper encoding of the genomic data can be learned and possibly used as a starting input of various inference tasks by combining this encoding with recent neural network-based tools for the reconstruction of recombination, demography or selection (Sheehan and Song 2016; Adrion et al. 2019; Flagel et al. 2019).

## Results

### Reconstructing genome wide population structure

Initially we created AGs with GAN, RBM, and two simple generative models for comparison: a Bernoulli and a Markov chain model (see Materials & Methods) using 2504 individuals (5008 haplotypes) from 1000 Genomes data (1000 Genomes Project Consortium et al. 2015), spanning 805 SNPs from all chromosomes which reflect a high proportion of the population structure present in the whole dataset (Colonna et al. 2014). Both GAN and RBM models capture a good portion of the population structure present in 1000 Genomes data while the other two models could only produce instances centered around 0 on principal component analysis (PCA) space (Figure 1). All major modes, corresponding to African, European and Asian genomes, are well represented in AGs produced by GAN and RBM models. Uniform manifold approximation and projection (UMAP) mapping results also correlate with the performed PCA (Supplementary Figure 3). We additionally checked the distribution of pairwise differences of haploid genomes to see how different AGs are from real genomes (Supplementary Figure 4). Both RBM and GAN models have highly similar distributions to the distribution of pairwise differences of the real genomes within themselves. Especially RBM excels at replicating the real peaks, indicating a high similarity with real genomes. Since GANs and RBMs showed an excellent performance for this use case, we further explored other characteristics using only these two models.

**Figure 1.**
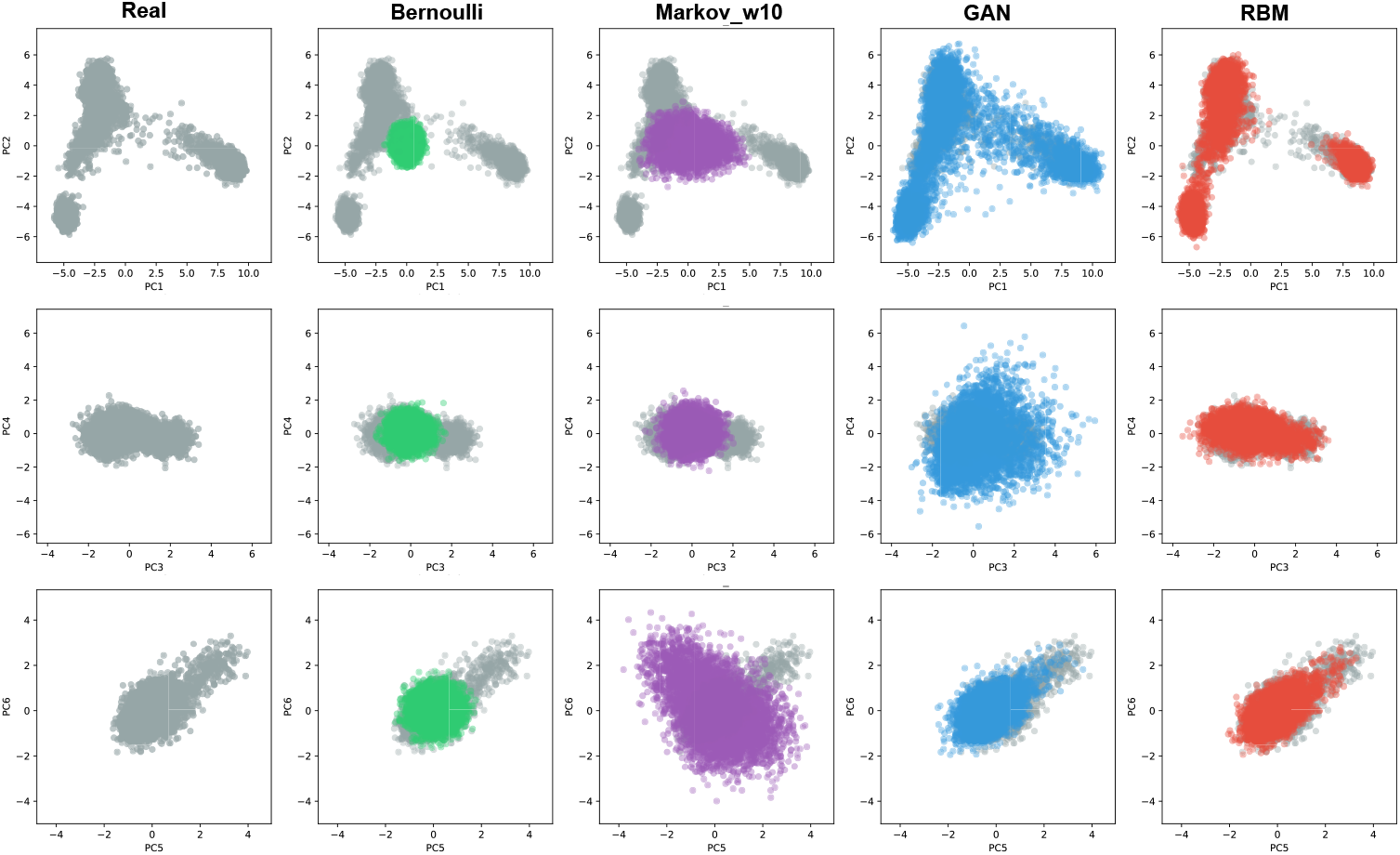
The six first axes of a PCA applied to real (gray) and artificial genomes (AGs) generated via Bernoulli (green), Markov chain (purple), GAN (blue) and RBM (red) models. There are 5000 haplotypes for each AG dataset and 5008 (2504 genomes) for the real dataset from 1000 Genomes spanning 805 informative SNPs. See Materials & Methods for detailed explanation of the generation procedures.

Furthermore, similarly to tSNE and UMAP, RBMs perform a non-linear dimension reduction of the data and provides a suitable representation of a genomic dataset as a by-product based on the non-linear feature space associated to the hidden layer (Supplementary Text). As Diaz-Papkovich et al (Diaz-Papkovich et al. 2019), we found that the RBM representation differs from the linear PCA ones. Here we plot the representation corresponding to the selected RBM model and exhibit its rapid evolution through training (Supplementary Figure 5).

Supplementary Figure 5 shows that African, East Asian, and to a lesser extent, European populations stand out on the two first components. The Finnish are slightly isolated from the other European (similar to Peruvian from American) populations on the first two components. South Asians are located at the center separated from Europeans, partially overlapping with American populations, and stand out at dimension 5 and higher. Interestingly when screening the hidden node activations, we observed that different populations or groups activate different hidden nodes, each one representing a specific combination of SNPs, thereby confirming that the hidden layer provides a meaningful encoding of the data (Supplementary Figure 6).

### Reconstructing local high-density haplotype structure

To evaluate if high quality artificial dense genome sequences can also be created by generative models, we applied the GAN and RBM methods to a 10K SNP region using (i) the same individuals from 1000 Genomes data and (ii) 1000 Estonian individuals from the high coverage Estonian Biobank (Leitsalu et al. 2015) to generate artificial genomes. PCA results of AGs spanning consecutive 10K SNPs show that both GAN and RBM models can still capture the relatively toned-down population structure (Supplementary Figure 7) as well as the overall distribution of pairwise distances (Supplementary Figure 8). Looking at the allele frequency comparison between real and artificial genomes, we see that especially GAN performs poorly for low frequency alleles, due to a lack of representation of these alleles in the AGs (Supplementary Figure 9). On the other hand, the distribution of the distance of real genomes to the closest AG neighbour shows that GAN model, although slightly underfitting, outperforms RBM model, for which an excess of small distances points towards slight overfitting (Supplementary Figure 10).

Additionally, we performed linkage disequilibrium (LD) analyses comparing artificial and real genomes to assess how successfully the AGs imitate short and long range correlations between SNPs. Pairwise LD matrices for real and artificial genomes all show a similar block pattern demonstrating that GAN and RBM accurately captured the overall structure with SNPs having higher linkage in specific regions (Figure 2a). However, plotting LD as a function of the SNP distance showed that all models capture weaker correlation, with RBM outperforming the GAN model perhaps due to its slightly overfitting characteristic (Figure 2b). To further determine the haplotypic integrity of AGs, we performed ChromoPainter (Lawson et al. 2012) and Haplostrips (Marnetto and Huerta-Sánchez 2017) analyses on AGs created using Estonians as the training data. It was visually impossible to distinguish the difference between real and artificial genomes in terms of local haplotypic structure with Haplostrips (Supplementary Figure 11). However, majority of the AGs produced via GAN model displayed an excess of short chunks when painted against 1000 Genomes individuals, whereas RBM AGs were nearly indistinguishable from real genomes (Supplementary Figure 12).

**Figure 2.**
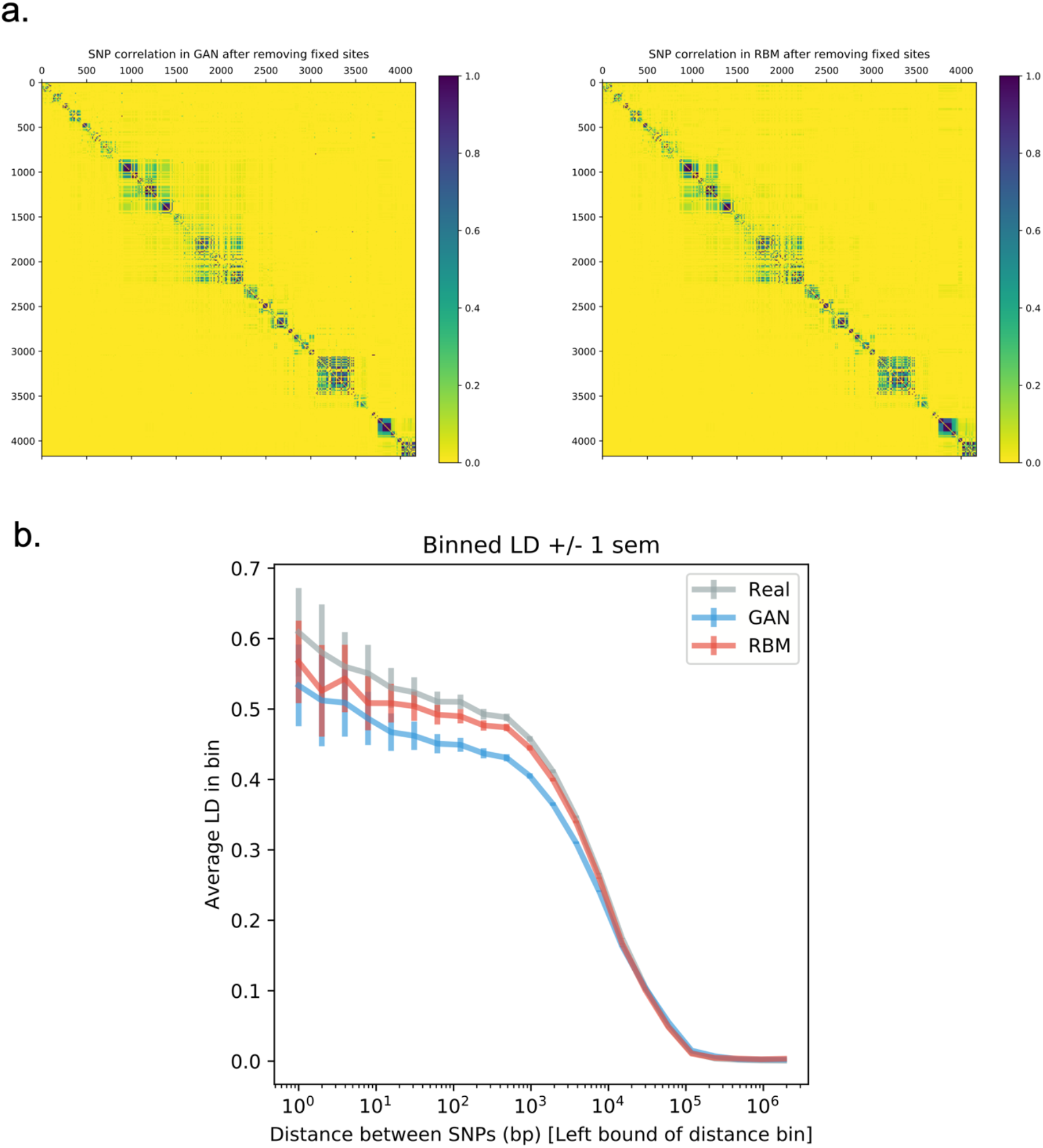
Linkage disequilibrium (LD) analysis on real and artificial Estonian genomes. **a)** Correlation (r^2^) matrices of SNPs. Lower triangular parts are SNP pairwise correlation in real genomes and upper triangular parts are SNP pairwise correlation in artificial genomes. **b)** LD as a function of SNP distance. Pairwise SNP distances were stratified into 50 bins and for each distance bin, the correlation was averaged over all pairs of SNPs belonging to the bin.

After demonstrating that our models generated realistic AGs according to the described summary statistics, we investigated further whether they respected privacy by measuring the extent of overfitting. We calculated two metrics of resemblance and privacy, the nearest neighbour adversarial accuracy (AA_TS_) and privacy loss presented in a recent study (Yale et al. 2019). AA_TS_ score measures whether two datasets were generated by the same distribution based on the distances between all data points and their nearest neighbours in each set. When applied to artificial and real datasets, a score between 0.5 and 1 indicates underfitting, between 0 and 0.5 overfitting (and likely privacy loss), and exactly 0.5 indicates that the datasets are indistinguishable. By using an additional real test set, it is also possible to calculate a privacy loss score that is positive in case of information leakage, negative otherwise, and approximately ranges from −0.5 to 0.5. Computed on our generated data, both scores support haplotypic pairwise difference results confirming the underfitting nature of GAN AGs and slightly overfitting nature of RBM AGs with a small risk of privacy leakage for the latter (Supplementary Figure 13).

Since it has been shown in previous studies that imputation scores can be improved using additional population specific reference panels (Gurdasani et al. 2015; Mitt et al. 2017), as a possible future use case, we tried imputing real Estonian genomes using 1000 Genomes reference panel and additional artificial reference panels with Impute2 software (Howie et al. 2011). Both combined RBM AG and combined GAN AG panels outperformed 1000 Genomes panel for the lowest MAF bin (for MAF < 0.05, 0.015 and 0.024 improvement respectively) which had 5926 SNPs out of 9230 total (Figure 3). Also mean info metric over all SNPs was 0.009 and 0.015 higher for combined RBM and GAN panels respectively, compared to the panel with only 1000 Genomes samples. However, aside from the lowest MAF bin, 1000 Genomes panel outperformed both concatenated panels for all the higher bins. This might be a manifestation of haplotypic deformities in AGs that might have disrupted the imputation algorithm.

**Figure 3.**
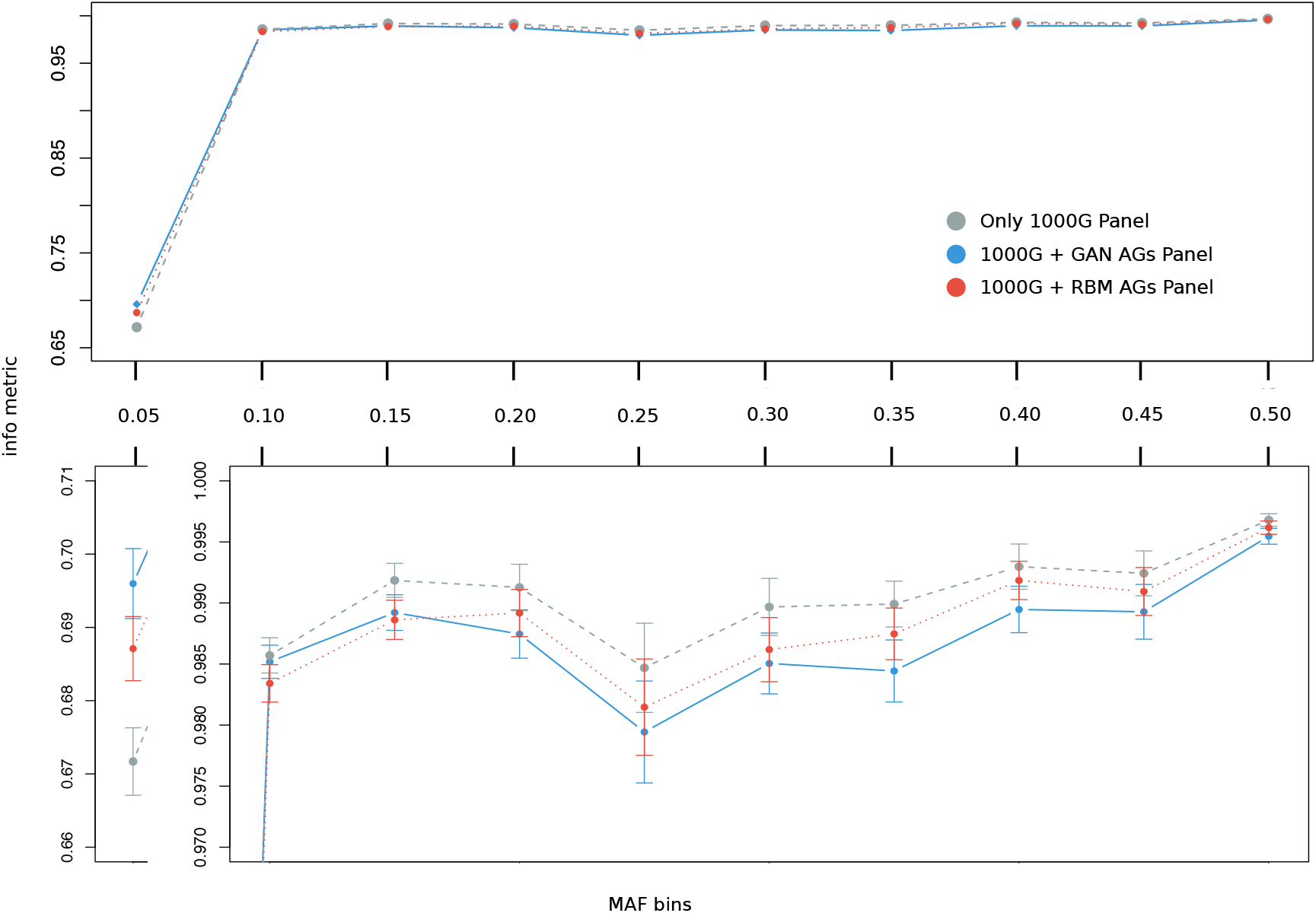
Imputation evaluation of three different reference panels based on Impute2 software’s info metric. Imputation was performed on 8678 Estonian individuals (which were not used in training of GAN and RBM models) using only 1000 Genomes panel (gray), combined 1000 Genomes and GAN artificial genomes panel (blue) and combined 1000 Genomes and RBM artificial genomes panel (red). SNPs were divided into 10 MAF bins, from 0.05 to 0.5, after which mean info metric values were calculated. Bars in the zoomed section show the standard error of mean.

### Selection tests

We additionally performed cross population extended haplotype homozygosity (XP-EHH) and population branch statistic (PBS) on a 3348 SNP region homogenously dispersed over chromosome 15 to assess if AGs can also be used for selection tests. Both XP-EHH and PBS results provided high correlation between the scores of real and artificial genomes (Figure 4). The peaks observed in real genome scores which might indicate possible selection signals were successfully captured by AGs.

**Figure 4.**
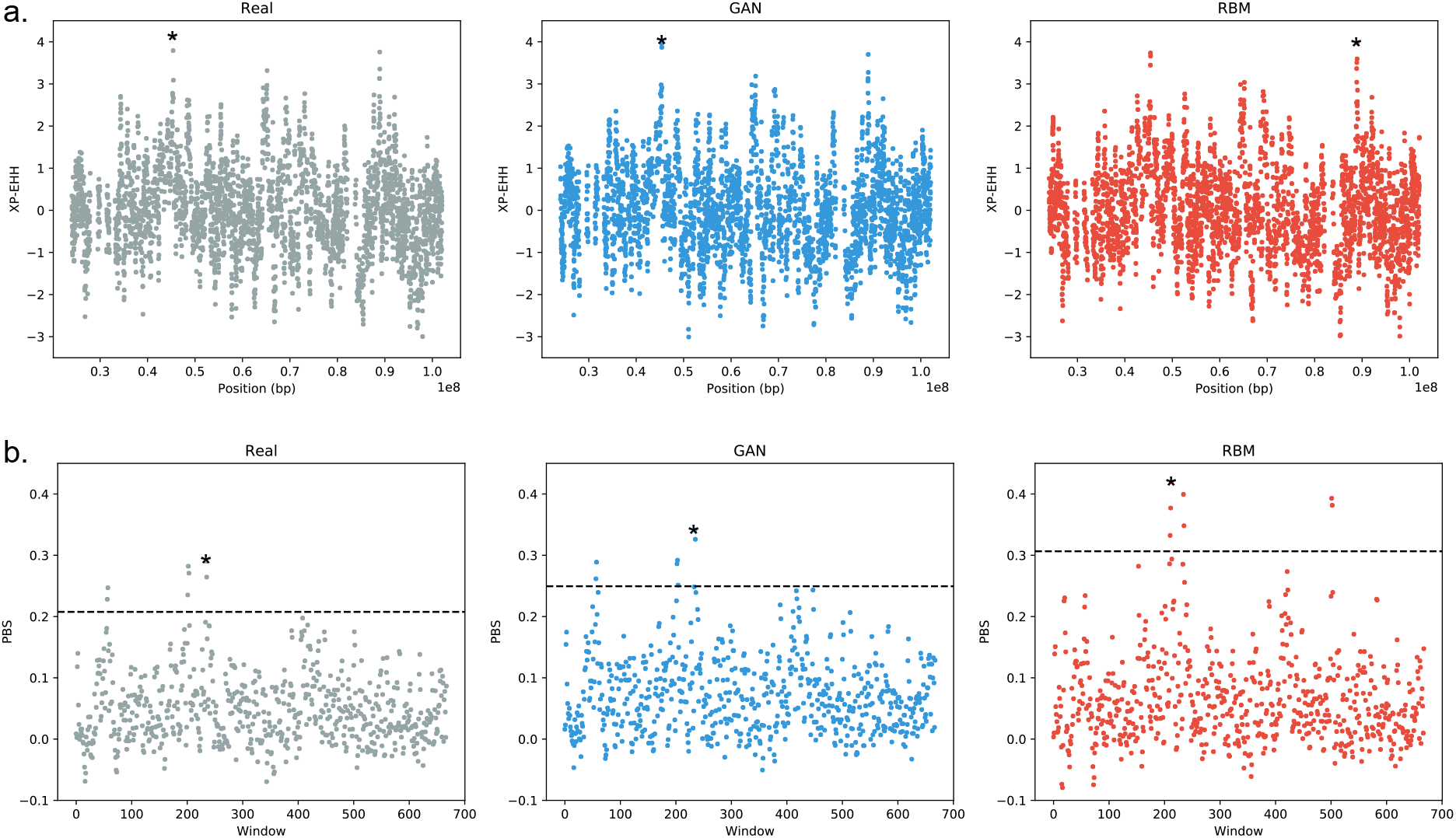
Selection tests on chromosome 15. **a)** Standardized XP-EHH scores of real and artificial Estonian genomes using 1000 Genomes Yoruba population (YRI) as the complementary population. Correlation coefficient between real and GAN XP-EHH score is 0.902, between real and RBM XP-EHH score is 0.887. **b)** PBS scores of real and artificial Estonian genomes using 1000 Genomes Yoruba (YRI) and Japanese (JPT) populations as the complementary populations. PBS window size is 10 and step size is 5. Dotted black line corresponds to the 99^th^ percentile. Correlation coefficient between real and GAN PBS score is 0.923, between real and RBM PBS score is 0.755. Highest peaks are marked by an asterisk.

### Linking genotypes with phenotypes

We then explored the possibility of creating AGs with unphased genotype data and recreating phenotype-genotype associations using generative models. As a proof of concept, we created GAN AGs via training on 1925 Estonian individuals with 5000 SNPs using unphased genotypes instead of haplotypes. There was an additional column in this dataset representing eye color (blue or brown). This region encompasses rs12913832 SNP which is highly associated with eye color phenotype (Han et al. 2008; Eriksson et al. 2010; Zhang et al. 2013). In our real genome dataset, nearly 96% of the individuals possessing at least one ancestral allele (A) have brown eye color while this percentage is 80% in GAN AGs. Similarly, 97% of all blue-eyed real individuals and 88% of the artificial ones are homozygous for the derived allele (G). No blue-eyed individuals are homozygous for the ancestral alleles in the real dataset and only 9 individuals out of the 1925 GAN AGs were homozygous ancestral with blue eyes (Supplementary Table). Chi-square tests based on the contingency tables were highly significant for both the real and artificial datasets (p-values < 2.2e-16 ; Supplementary Table 1). These results suggest that AGs were able to reproduce the genotypic-phenotypic association existing in the real dataset. We could not detect the same association in the RBM AG dataset (see Discussion).

## Discussion

In this study, we applied generative models to produce artificial genomes and evaluated their characteristics. To the best of our knowledge, this is the first application of GAN and RBM models in population genetics context, displaying overall promising applicability. We showed that population structure and frequency-based features of real populations can successfully be preserved in AGs created using GAN and RBM models. Furthermore, both models can be applied to sparse or dense SNP data given a large enough number of training individuals. Our different trials showed that the minimum required number of individuals for training is highly variable, possibly correlated with the diversity among individuals (data not shown). Since haplotype data is more informative, we created haplotypes for the analyses but we also demonstrated that the GAN model can be applied to genotype data too, by simply combining two haplotypes if the training data is not phased (see Materials & Methods). In addition, we showed that it is possible to generate AGs with simple phenotypic traits through genotype data (see Results). Even though there were only two simple classes, blue and brown eye color phenotypes, generative models can be improved in the future to hold the capability to produce artificial datasets combining AGs with multiple phenotypes. The training of the RBM in this case did not work properly. We believe that it is because the encoding of the phenotype is not well-suited for the RBM. Further investigation on that part would be out of the scope of this article, but we suspect that an encoding of the type “one-hot” vector addition of a stronger learning rate for the weights linked to the phenotype nodes could improve the training.

One major drawback of the proposed models is that, due to computational limitations, they cannot yet be harnessed to create whole artificial genomes but rather snippets or sequential dense chunks. Although parallel computing might be a solution, this might further disrupt the haplotype structure in AGs. Instead, adapting convolutional GANs for AG generation might be a possible solution in the future (Radford et al. 2016). Another problem arose due to rare alleles, especially for the GAN model. We showed that nearly half of the alleles become fixed in the GAN AGs in the 10K SNP dataset, whereas RBM AGs capture more of the rare alleles present in real genomes (Supplementary Figure 14). A known issue in GAN training is mode collapse (Salimans et al. 2016), which occurs when the generator fails to cover the full support of the data distribution. This failure case could explain the inability of GANs to generate rare alleles. For some applications relying on rare alleles, GAN models less sensitive to mode dropping would be a promising alternative (Arjovsky et al. 2017; Lucas et al. 2018).

An important use case for the AGs in the future might be creating publicly available versions of private genome banks. Through enhancements in scientific knowledge and technology, genetic data becomes more and more sensitive in terms of privacy. AGs might offer a versatile solution to this delicate issue in the future by protecting the anonymity of real individuals. Our results showed that GAN AGs are possibly underfitting while, on the contrary, RBM AGs are slightly overfitting, based on distribution of minimum distance to the closest neighbour (Supplementary Figure 10) and AA_TS_ scores (Supplementary Figure 13a), although we showed how overfitting could be restrained by integrating AA_TS_ scores within our models as a criterion for early stopping in training (before the networks start overfitting). In the context of the privacy issue, GAN AGs have a slight advantage since underfitting is preferable. More distant AGs would hypothetically be harder to be traced back to the original genomes. We also tested the sensitivity of the AA_TS_ score and privacy loss (Supplementary Figure 15). It appears that both scores are affected very slightly when we add only a few real genomes to the AG dataset from the training set. Although this case is easily detectable by examining the extreme left tail of the pairwise distribution, it advocates for combining multiple privacy loss criteria and developing other sensitive measurement techniques for better assessment of generated AGs. Additionally, even though we did not detect exact copies of real genomes in AG sets created either by RBM or GAN models, it is a very complicated task to determine if the generated samples can be traced back to the originals. Reliable measurements need to be developed in the future to assure complete anonymity of the source individuals given the released AGs. In particular, we will investigate whether the differential privacy framework is performant in the context of large population genomics datasets (Dwork et al. 2006; Torkzadehmahani et al. 2019).

Imputation results demonstrated promising outcomes especially for population specific low frequency alleles. However, imputation with both RBM and GAN AGs integrated reference panels showed slight decrease of info metric for higher frequency alleles compared to only 1000 Genomes panel (Figure 3). We initially speculated that this might be related to the disturbance in haplotypic structure and therefore, tried to filter AGs based on chunk counts from ChromoPainter results, preserving only AGs which are below the average chunk count of real genomes. The reasoning behind this was to preserve most real-alike AGs with undisturbed chunks. Even with this filtering, slight decrease in higher MAF bins was still present (data not shown). Yet results of implementation with AGs for low frequency alleles and without AGs for high frequency ones could be combined to achieve best performance. In terms of imputation, future improved models can become practically very useful, largely for GWAS studies in which imputation is a common application to increase resolution. Different generative models such as MaskGAN (Fedus et al. 2018) which demonstrated good results in text gap filling might also be adapted for genetic imputation. RBM is possibly another option to be used as an imputation tool directly by itself, since once the weights have been learned, it is possible to fix a subset of the visible variables and to compute the average values of the unobserved ones by sampling the probability distribution (in fact, it is even easier than sampling entirely new configurations since the fixed subset of variables will accelerate the convergence of the sampling algorithm).

Scans for detecting selection are another promising use case for AGs. The XP-EHH and PBS scores computed on AGs were highly correlated with the scores of real genomes. In particular, the highest peak we obtained for Estonian genomes was also present in AGs, although it was the second highest peak in RBM XP-EHH plot (Figure 4). This peak falls within the range of skin color associated *SLC24A5* gene, which is potentially under positive selection in many European populations (Basu Mallick et al. 2013).

As an additional feature, training an RBM to model the data distribution gives access to a latent encoding of data points, providing a potentially easier to use representation of data (Supplementary Figure 5). Future works could augment our current GAN model to also provide an encoding mechanism, in the spirit of (Dumoulin et al. 2016 Jun 2), (Chen et al. 2016) or (Donahue et al. 2016). These interpretable representations of the data are expected to be more relevant for downstream tasks (Chen et al. 2016) and could be used as a starting point for various population genetics analyses such as demographic and selection inference, or yet unknown tasks.

Although there are some current limitations, generative models will most likely become prominent for genetics in the near future with many promising applications. In this work, we demonstrated the first possible implementations and use of AGs in the forthcoming field which we would like to name artificial genomics.

## Materials & Methods

### Data

We used 2504 individual genomes from 1000 Genomes Project (1000 Genomes Project Consortium 2015) and 1000 individuals from Estonian Biobank (Leitsalu et al. 2015) to create artificial genomes (AGs). Additional 2000 Estonians were used as a test dataset. Another Estonian dataset consisting of 8678 individuals which were not used in training were used for imputation. Analyses were applied to a highly differentiated 805 SNPs selected as a subset from (Colonna et al. 2014), 3348 SNPs dispersed over the whole chromosome 15 and a dense 10000 SNP range/region from chromosome 15. We also used a narrowed down version of the same region from chromosome 15 with 5000 SNPs with an additional eye color column for unphased genotype data using another 1925 Estonians as training dataset. In this set, 958 of the Estonian samples have brown (encoded as 1) and 967 have blue eyes (encoded as 0). In the data format we used, rows are individuals/haplotypes (instances) and columns are positions/SNPs (features). Each allele at each position is represented either by 0 or 1. In the case of phased data (haplotypes), each column is one position whereas in the case of unphased data, each two column corresponds to a single position with alleles from two chromosomes.

### GAN Model

We used python-3.6, Keras 2.2.4 deep learning library with TensorFlow backend (Chollet 2015), pandas 0.23.4 (McKinney 2010) and numpy 1.16.4 (Oliphant 2007) for the GAN code. Generator of the GAN model we present consists of an input layer with the size of the latent vector size 600, one hidden layer with size proportional to the number of SNPs as SNP_number/1.2 rounded, another hidden layer with size proportional to the number of SNPs as SNP_number/1.1 rounded and an output layer with the size of the number of SNPs. The latent vector was set with numpy.random.normal function setting the mean of the distribution as 0 and the standard deviation as 1. The discriminator consists of an input layer with the size of the number of SNPs, one hidden layer with size proportional to the number of SNPs as SNP_number/2 rounded, another hidden layer with size proportional to the number of SNPs as SNP_number /3 rounded and an output layer of size 1. All layer outputs except for output layers have LeakyReLU activation functions with leaky_alpha parameter 0.01 and L2 regularization parameter 0.0001. The generator output layer activation function is tanh and discriminator output layer activation function is sigmoid. Both discriminator and combined GAN were compiled with Adam optimization algorithm with binary cross entropy loss function. We set the discriminator learning rate as 0.0008 and combined GAN learning rate as 0.0001. For 5000 SNP data, the discriminator learning rate was 0.00008 and combined GAN learning rate was 0.00001. Training to test dataset ratio was 3:1. We used batch size of 32 and trained all datasets up to 20000 epochs. We tried stopping the training based on AA_TS_ scores. The score was calculated at 200 epoch intervals. For 805 SNP data, AA_TS_ converged very quickly close to optimum 0.5 score. However, the difference between AA_truth_ and AA_syn_ scores indicates possible overfitting to multiple data points so it was difficult to define a stopping point. For 10K SNP data, convergence was observed after ~30K epochs (to around 0.75) and reduced the number of fixed alleles in AGs but the gain was very minimal (Supplementary Figure 16). Additionally, GAN was prone to mode collapse especially after 20K epochs which resulted in multiple failed training attempts. Therefore, we stopped training based on coherent PCA results of AGs with real genomes. During each batch in the training, when only the discriminator is trained, we applied smoothing to the real labels (1) by vectoral addition of random uniform distribution via numpy.random.uniform with lower bound 0 and upper bound 0.1. Elements of the generated outputs were rounded to 0 or 1 with numpy.rint function.

### RBM Model

The RBM was coded in Julia (Bezanson et al. 2017), and all the algorithm for the training has been done by the authors. The part of the algorithm involving linear algebra used the standard package provided by Julia. Two versions of the RBM were considered. In both versions, the visible nodes were encoded using Bernoulli random variables {0,1}, and the size of the visible layer was the same size as the considered input. Two different types of hidden layers were considered. First with a sigmoid activation function (hence having discrete {0,1} hidden variables), second with ReLu (Rectified Linear unit) activations in which case the hidden variables were positive and continuous (there are distributed according to a truncated gaussian distribution when conditioning on the values of the visible variables). Results with sigmoid activation function were worse compared to ReLu so we used ReLu for all the analyses (Supplementary Figure 17MY). The number of hidden nodes considered for the experiment was Nh=100 for the 805 SNP dataset and Nh=500 for the 10k one. There is no canonical way of fixing the number of hidden nodes, in practice we checked that the number of eigenvalues learnt by the model was smaller than the number of hidden nodes, and that by adding more hidden nodes no improvement were observed during the learning. The learning in general is quite stable, in order to have a smooth learning curve, the learning rate was set between 0.001 and 0.0001 and we used batch size of 32. The negative term of the gradient of the likelihood function was approximated using the PCDk method (Brügge et al. 2013), with k=10 and 100 of persistent chains. As a stopping criterion, we looked at when the AA_TS_ score converges to the ideal value of 0.5 when sampling the learned distribution. When dealing with large and sparse datasets for selection tests, RBM model did not manage to provide reasonable AA_TS_ scores because the sampling is intrinsically difficult for large systems with strong correlation. In that case, we used coherent PCA results as a stopping criterion.

### Bernoulli Distribution Model

We used python-3.6, pandas 0.23.4 and numpy 1.16.4 for the Bernoulli distribution model code. Each allele at a given position was randomly drawn given the derived allele frequency in the real population.

### Markov Chain Model

We used python-3.6, pandas 0.23.4 and numpy 1.16.4 for the Markov chain model code. Allele at the initial position was set by drawing from a Bernoulli distribution parameterized with the real frequency. Each successive allele was then drawn randomly according to its probability given the previous sequence window. After the initial position, the sequence window size increased incrementally up to a predifined window size (5 or 10 SNPs).

### Chromosome Painting

We compared the haplotype sharing distribution between real and artificial chromosomes through ChromoPainter (Lawson et al. 2012). In detail, we have painted 100 randomly selected “real” and “artificial” Estonians (recipients) against all the 1000 Genome Project phased data (donors). The nuisance parameters -n (348.57) and -M (0.00027), were estimated running 10 iterations of the expectation-maximization algorithm on a subset of 3,800 donor haplotypes.

### Haplostrips

We used Haplostrips (Marnetto and Huerta-Sánchez 2017) to visualize the haplotype structure of real and artificial genomes. We extracted 500 individuals from each sample set (Real, GAN AGs, RBM AGs) and considered them as different populations. Black dots represent derived alleles, white dots represent ancestral alleles. The plotted SNPs were filtered for a population specific minor allele frequency >5%; haplotypes were clustered and sorted for distance against the consensus haplotype from the real set. See the application article for further details about the method.

### Nearest Neighbour Adversarial Accuracy (AA_TS_) and Privacy Loss

We used the following equations for calculating AA_TS_ and privacy loss scores (Yale et al. 2019):

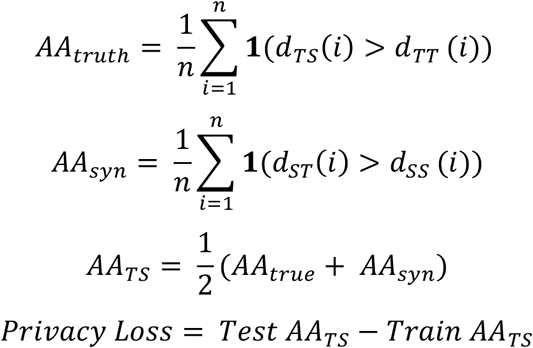

where *n* is the number of real samples as well as of artificial samples; 1 is a function which takes the value 1 if the argument is true and 0 if the argument is false; *d_TS_*(*i*) is the distance between the real genome indexed by i and its nearest neighbour in the artificial genome dataset; *d_ST_*(*i*) is the distance between the artificial genome indexed by i and its nearest neighbour in the real genome dataset; *d_TT_*(*i*) is the distance of the real genome indexed by i to its nearest neighbour in the real genome dataset; *d_SS_*(*i*) is the distance of the artificial genome indexed by i to its nearest neighbour in the artificial genome dataset. An AA_TS_ score of 0.5 is optimal whereas lower values indicate overfitting and higher values indicate underfitting. For a better resolution for the detection of overfitting, we also provided AA_truth_ and AA_syn_ metrics identified in the general equation of AA_TS_. If AA_TS_ ≈ 0.5 but AA_truth_ ≈ 0 and AA_syn_ ≈ 1, this means that the model is not overfitting in terms of a single data point but multiple ones. In other words, the model might be focusing on small batches of similar real genomes to create artificial genomes clustered at the center of each batch.

### Selection Tests

We used scikit-allel package for XP-EHH (Sabeti et al. 2007) and PBS (Yi et al. 2010) tests. We used 1000 Estonian individuals (2000 haplotypes) with 3348 SNPs which were homogenously dispersed over chromosome 15 for the training of GAN and RBM models. For XP-EHH, Yoruban (YRI, 216 haplotypes) population from 1000 Genomes data was used as the complementary population. For PBS, Yoruban (YRI, 216 haplotypes) and Japanese (JPT, 208 haplotypes) populations from 1000 Genomes data were used as complementary populations. PBS window size was 10 and step size was 5, resulting in 668 windows. 216 real and 216 AG haplotypes were compared for the analyses.

## Acknowledgements

This work was supported by the European Union through the European Regional Development Fund (Project No. 2014-2020.4.01.16-0024, MOBTT53: LP, DM, BY; Project No. 2014-2020.4.01.16-0030: LO, FM); the Estonian Research Council grant PUT (PRG243): LP; Laboratoire de Recherche en Informatique: FJ. Thanks to Inria TAU team for providing computational resources. Thanks to Adrien Pavao for his valuable insight into AA_TS_ score.

