## Supplementary Figure for "Creating Artificial Human Genomes Using Generative Models"

**Supplementary Figure 1.** Generative adversarial network (GAN) scheme.

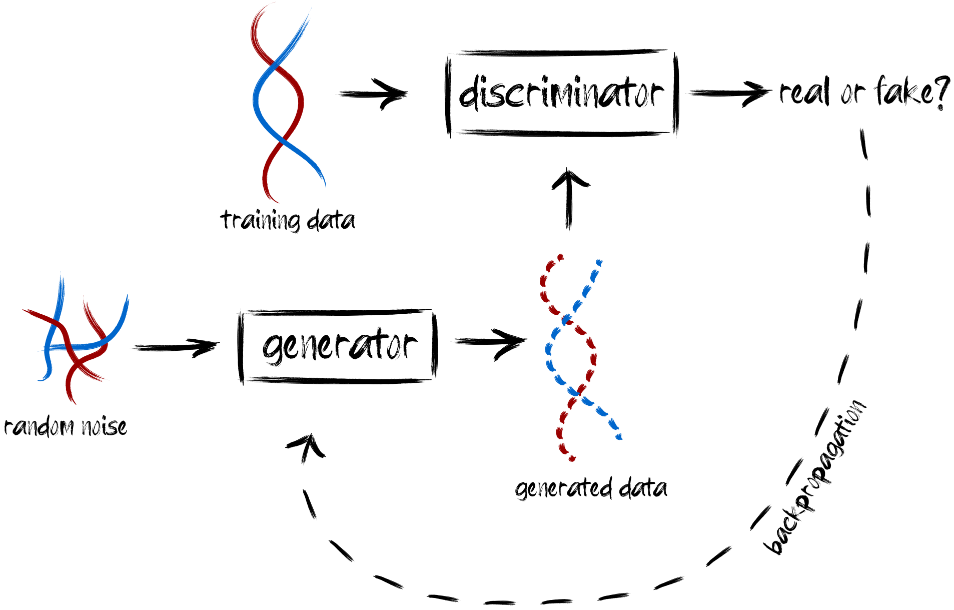

**Supplementary Figure 2.** Restricted Boltzmann machine (RBM) scheme.

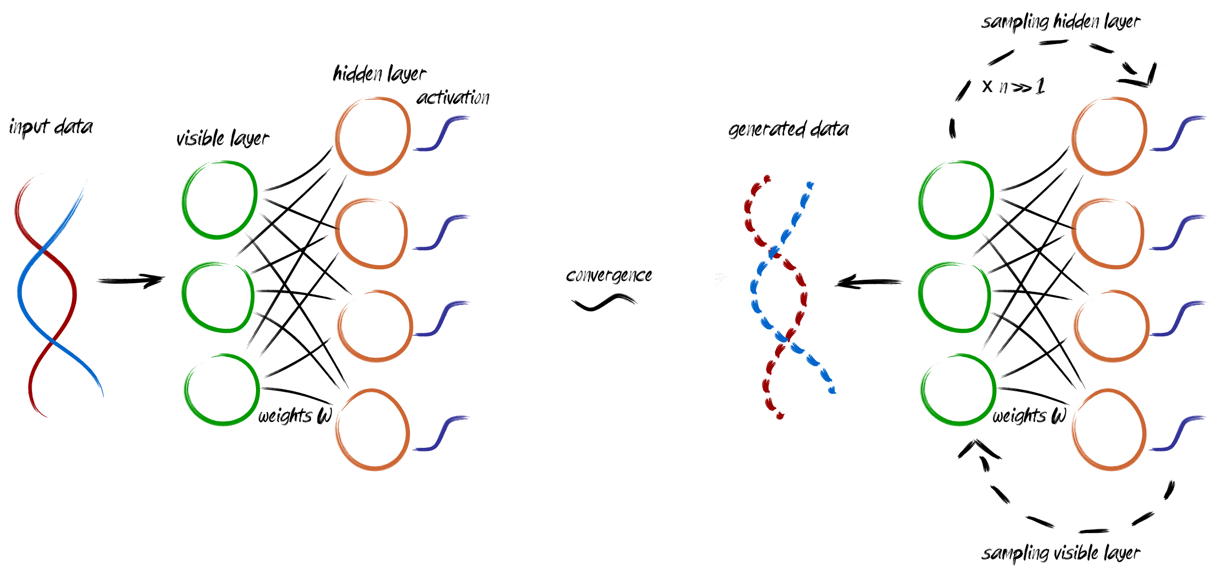

**Supplementary Figure 3.** Uniform manifold approximation and projection (UMAP) of real genomes from 1000 Genomes data spanning 805 SNPs along with artificial genome counterparts created via **a)** Bernoulli, **b)** Markov chain (with 10 window length), **c)** GAN and **d)** RBM models.

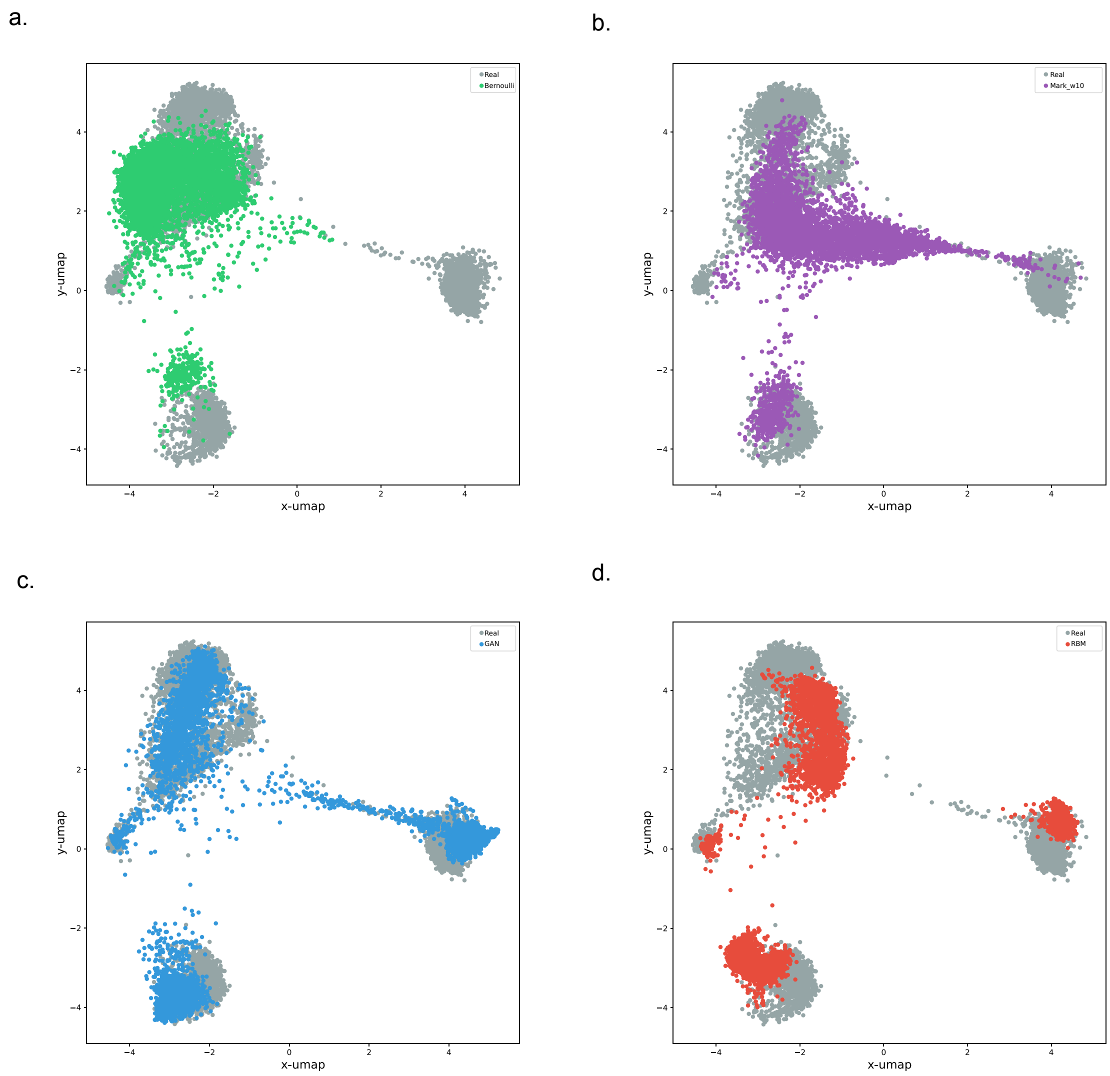

**Supplementary Figure 4.** Distribution of haplotypic pairwise difference within (left) and between (right) datasets of real genomes from 1000 Genomes data spanning 805 SNPs and artificial genome counterparts generated using different models.

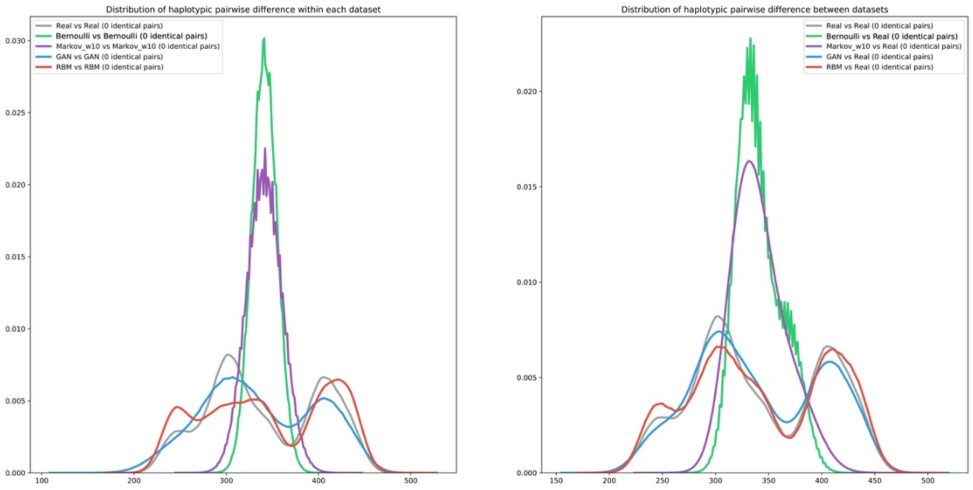

**Supplementary Figure 5.** Comparison of PCA (right column) and non-linear dimension reduction via RBM (left column) for real genomes from 1000 Genomes data spanning 805 SNPs. The RBM reduction was obtained by projecting the real data into the hidden space of the RBM (see Supplementary Text). Population codes are as defined by the 1000 Genomes Project.

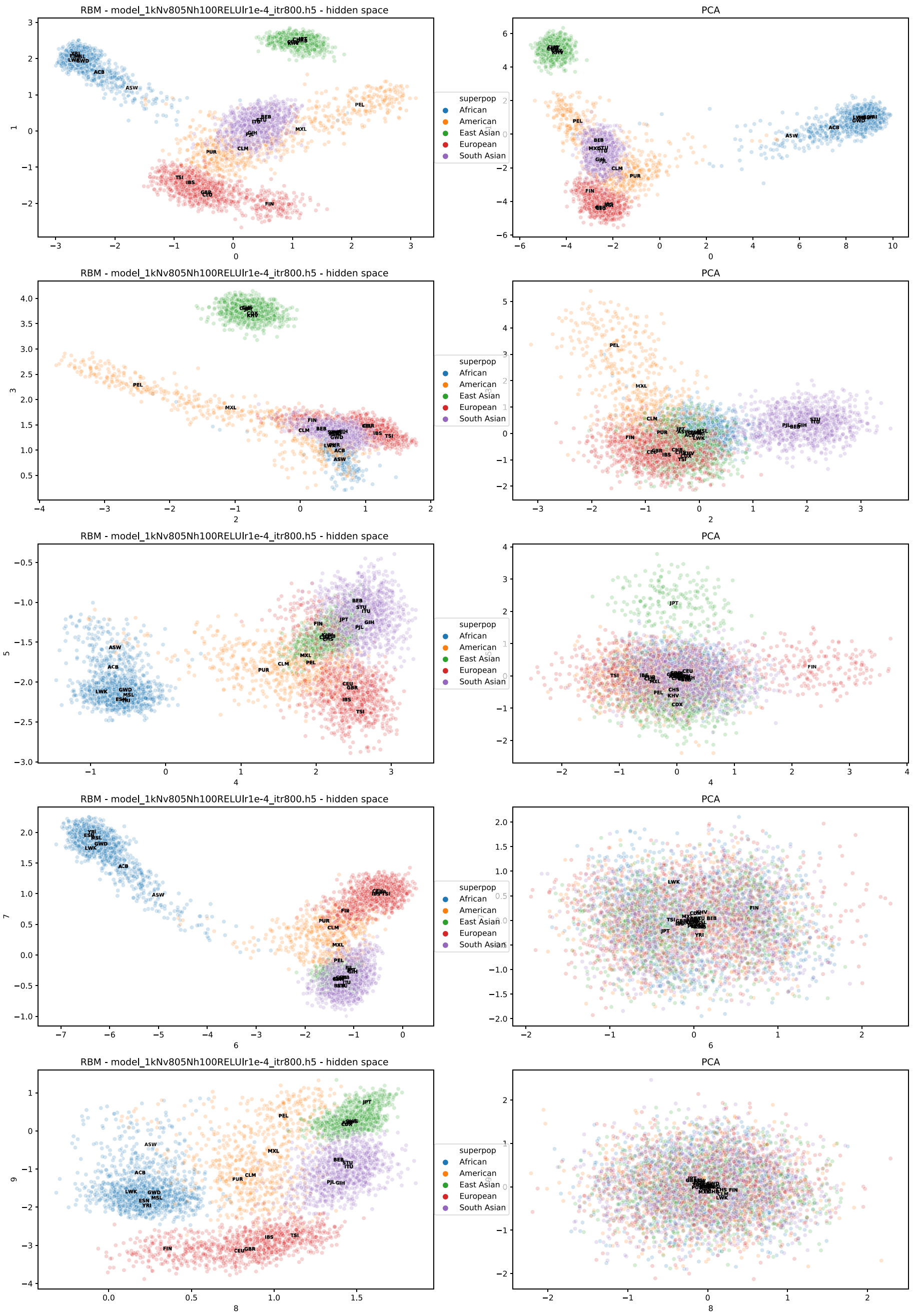

**Supplementary Figure 6.** Activations of each of the 100 nodes belonging to the RBM hidden layer when applied to the real genomes from 1000 Genomes data spanning 805 SNPs. For each hidden node the X-axis corresponds to the real haplotypes and Y-axis to the activation of the node by a single haplotype. On the X-axis, haplotypes are ordered by region (Africa, America, East Asia, European, East Asia) and colored by population. Because this RBM activation function is a ReLU with threshold 0 (by design), all values are positive and a zero-value indicates that the node is not activated by a given haplotype. The ordering of nodes has no specific meaning.

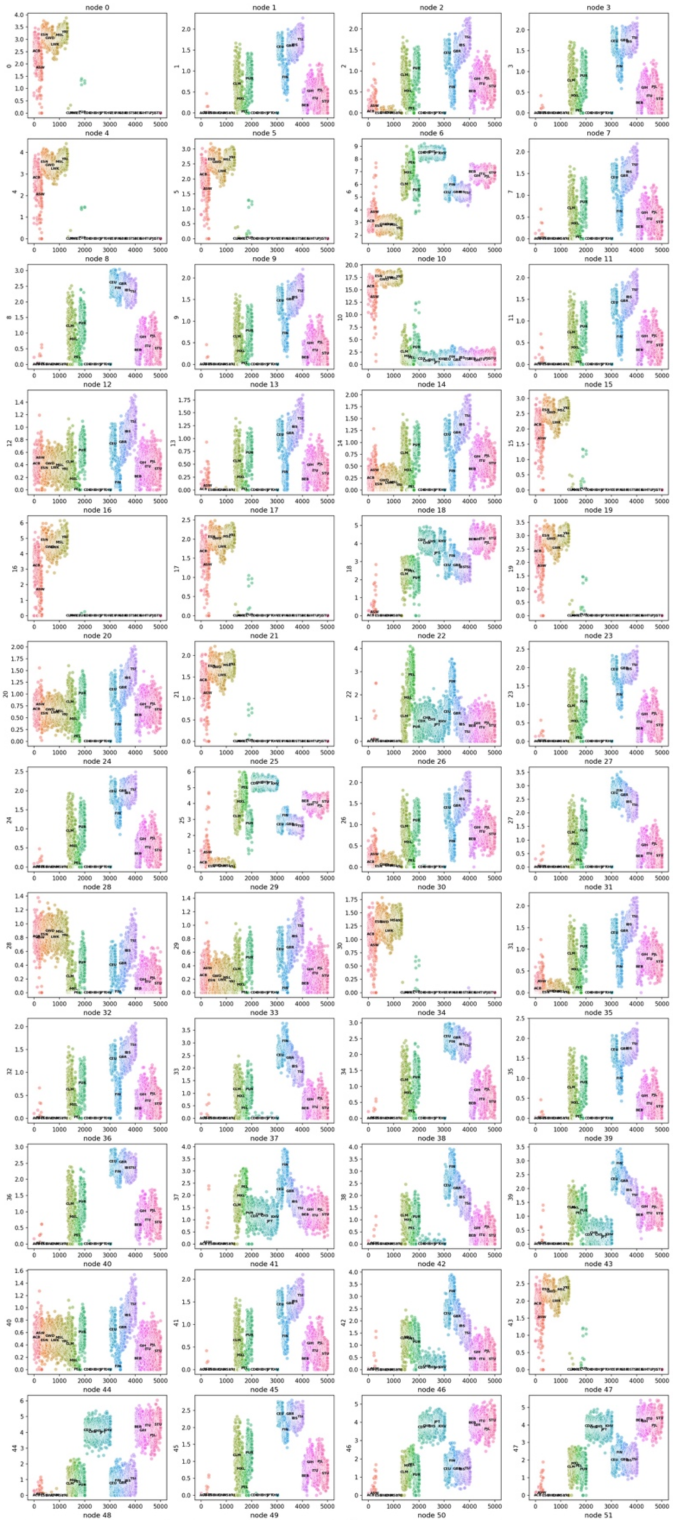

**Supplementary Figure 6 (Cont'd).**

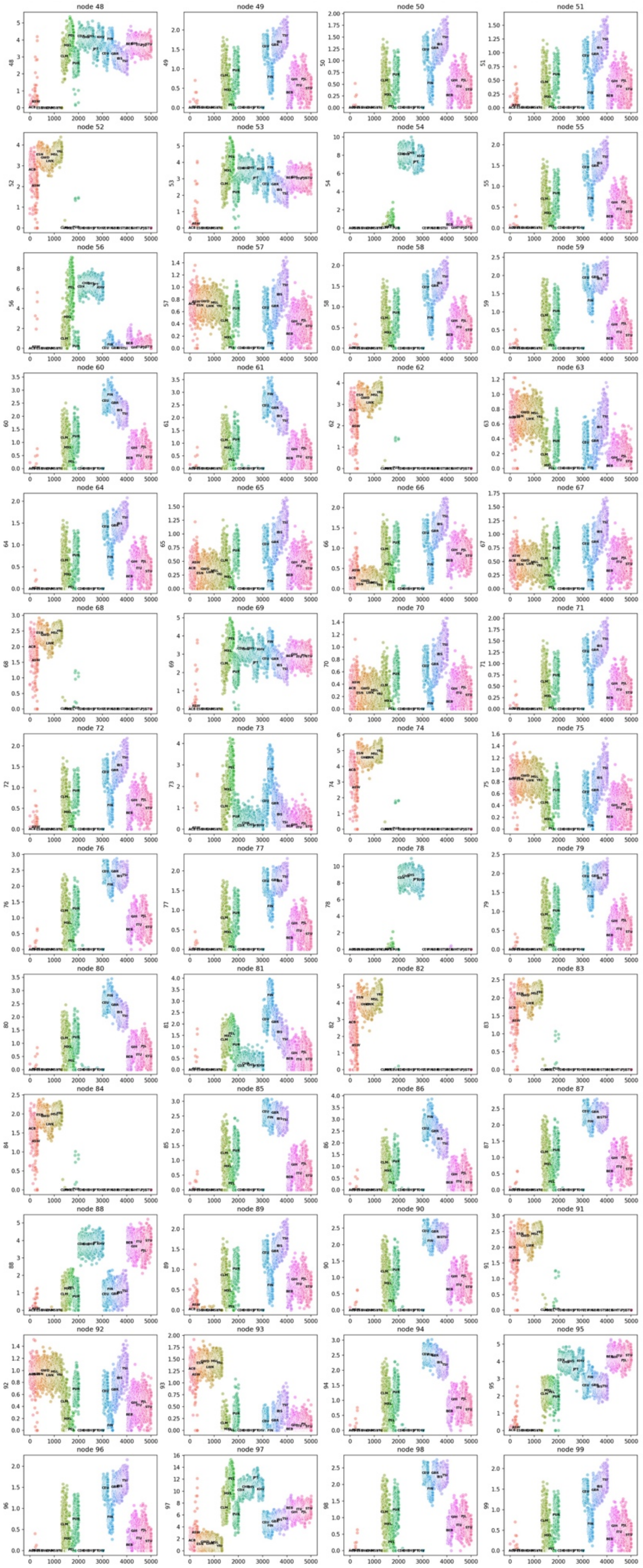

**Supplementary Figure 7.** PCA of real genomes (gray) from **a)** 1000 Genomes data and **b)** Estonian Biobank spanning 10K SNPs along with artificial genome counterparts generated using GAN (blue) and RBM (red) models.

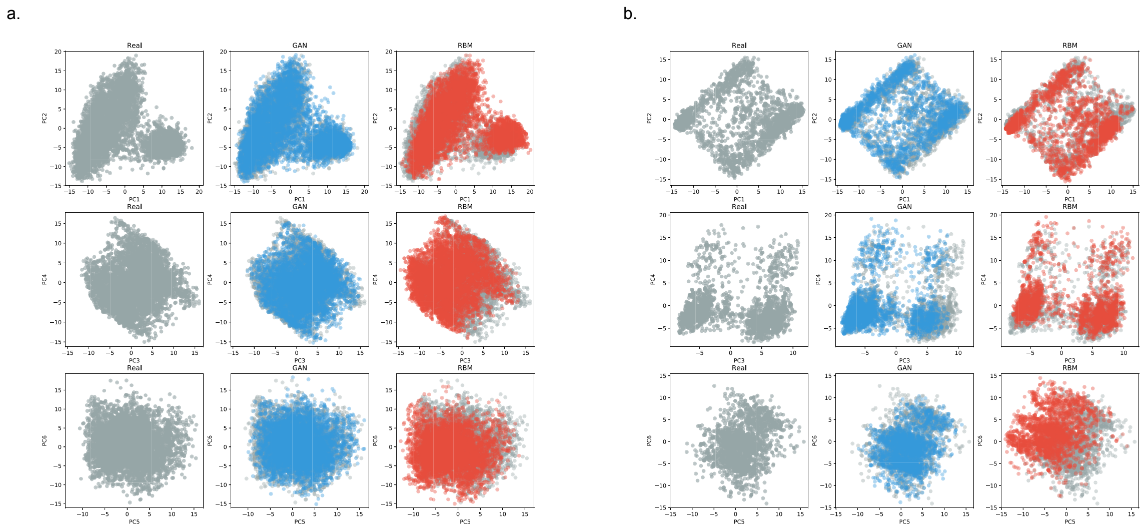

**Supplementary Figure 8.** Distribution of haplotypic pairwise difference within (left) and between (right) datasets of real genomes from **a)** 1000 Genomes data and **b)** Estonian Biobank spanning 10K SNPs and artificial genome counterparts generated using GAN and RBM models.

a.

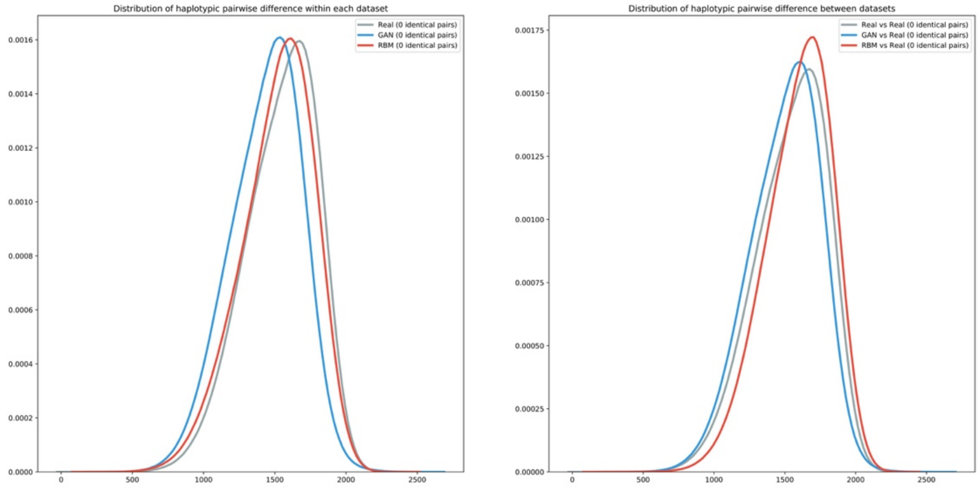


b.

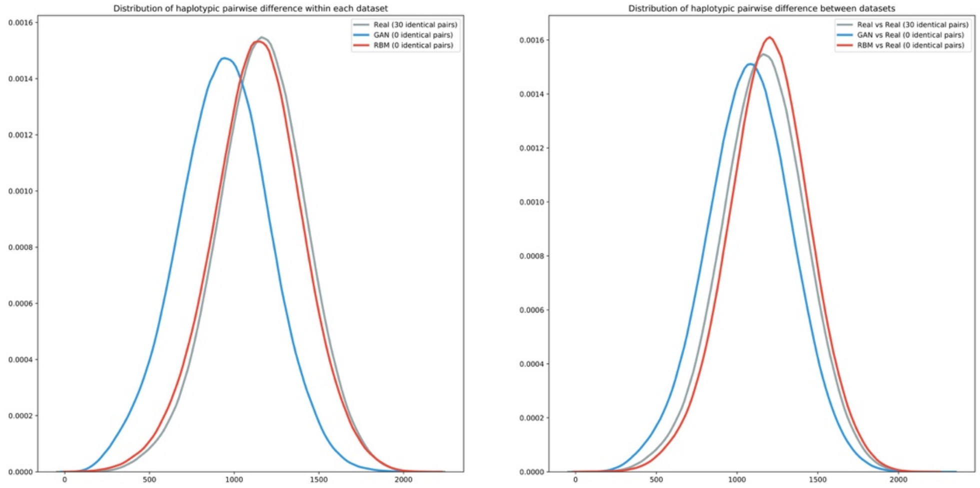

**Supplementary Figure 9.** Allele frequency comparison between real genomes from Estonian Biobank spanning 10K SNPs and artificial genome counterparts generated using GAN and RBM models as **a)** the whole range and **b)** zoomed to low frequencies. Clustering below the diagonal in the low frequency section for the GAN plot indicates insufficient representation of rare alleles in artificial genomes.

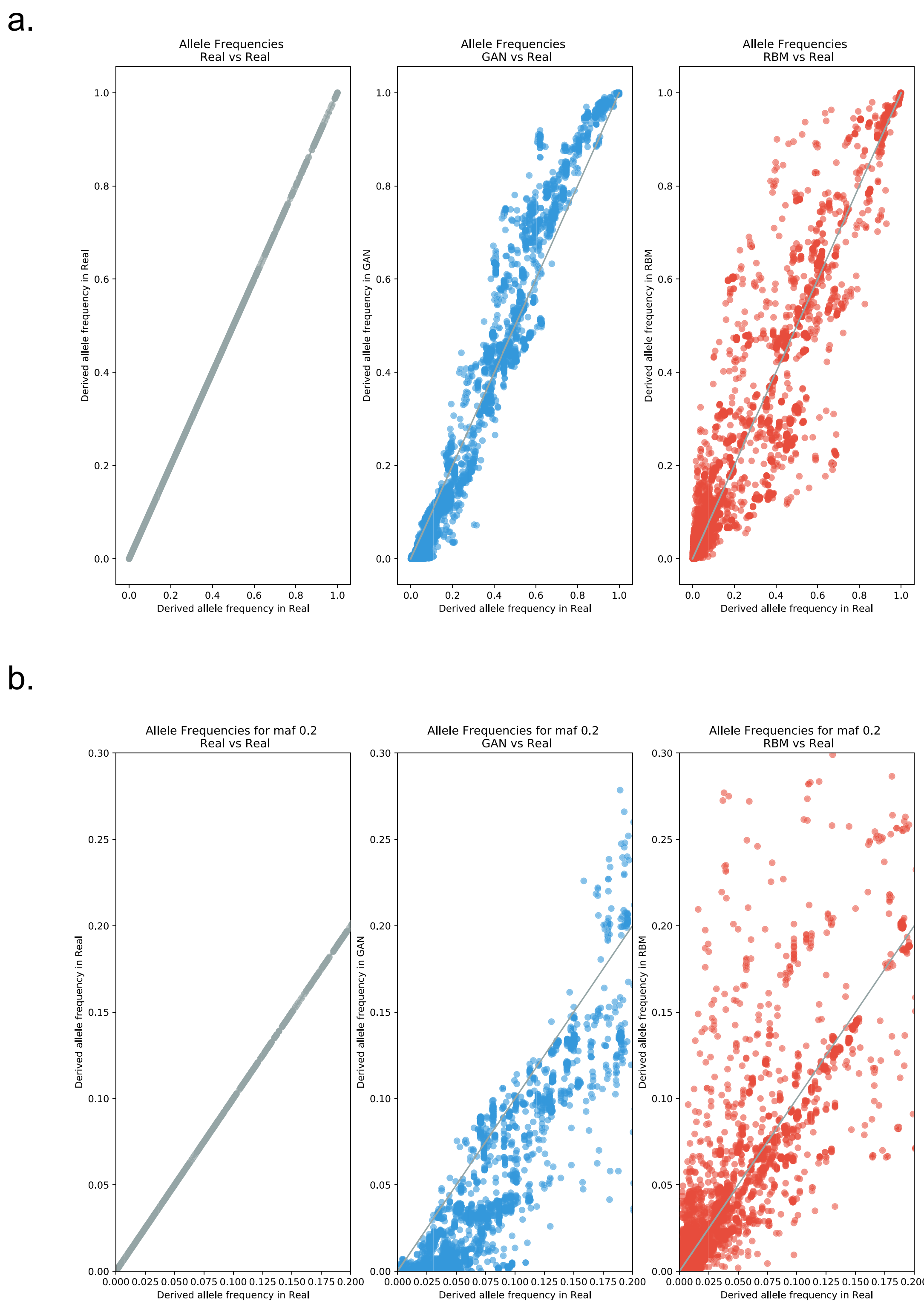

**Supplementary Figure 10.** Distribution of minimum distance to the closest neighbour for real genomes from **a)** 1000 Genomes data and **b)** Estonian Biobank spanning 10K SNPs along with artificial genome counterparts generated via GAN and RBM models.

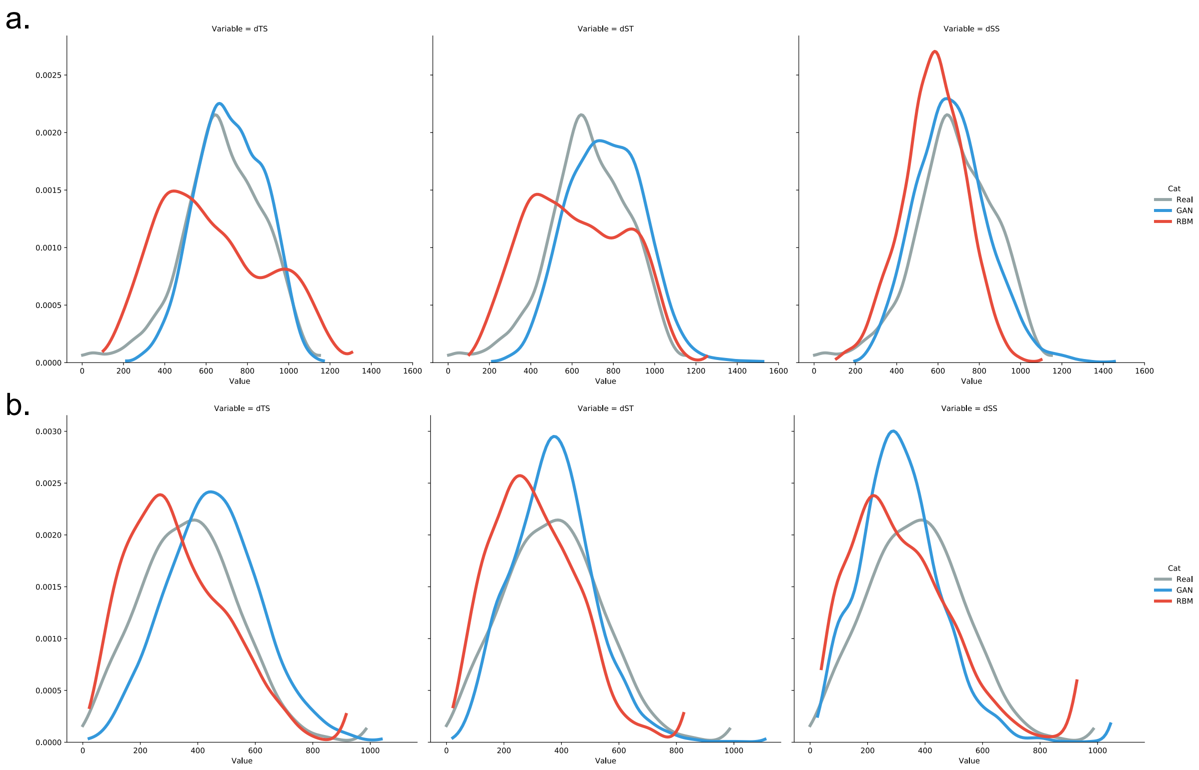

**Supplementary Figure 11.** Haplostrips showing the mixed nature of haplotype structures for real Estonian (gray rows) along with GAN (blue rows) and RBM (red rows) haplotypes.

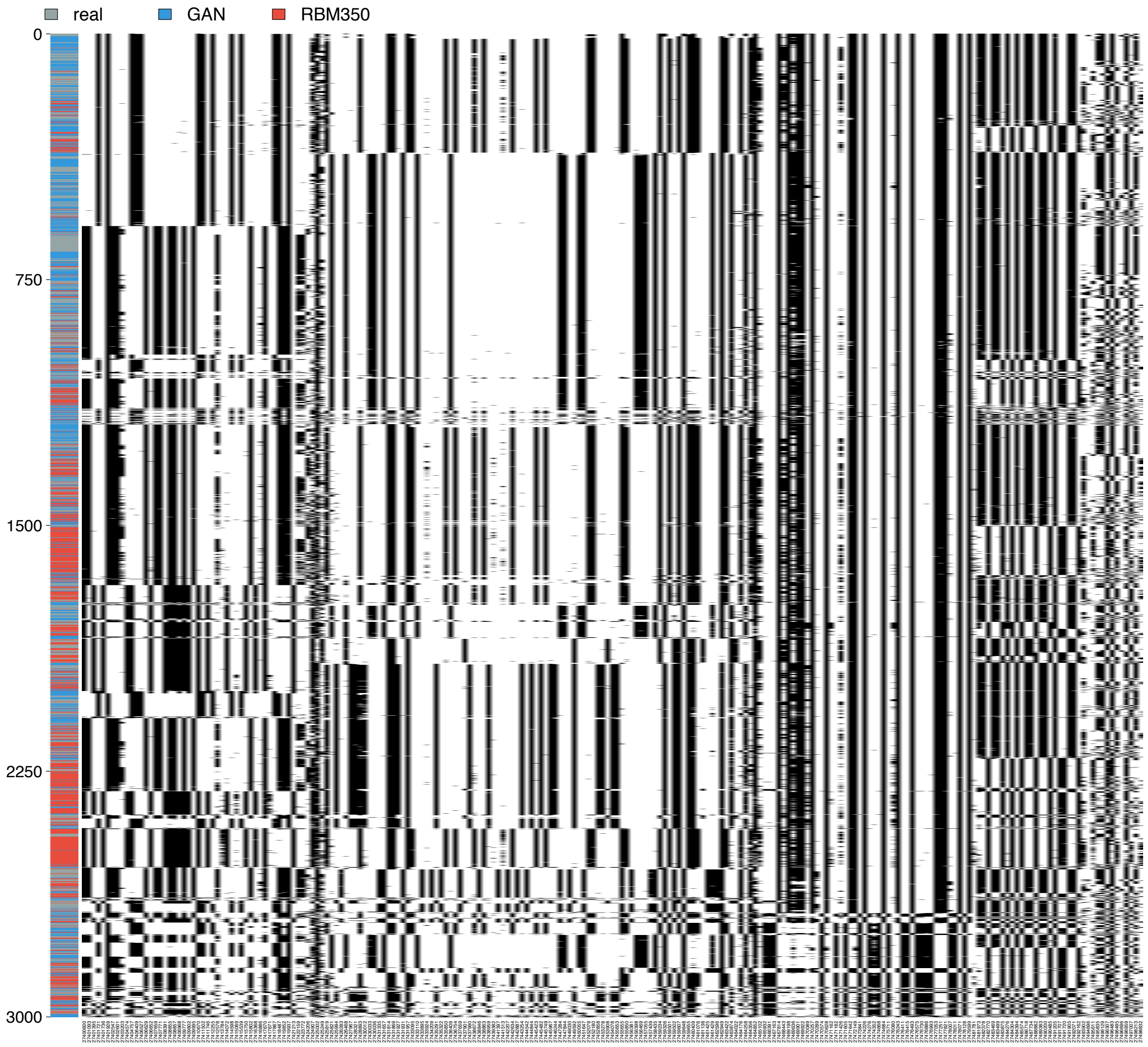

**Supplementary Figure 12.** Chromosome painting of **a)** real Estonian genomes, **b)** GAN and **c)** RBM artificial Estonian genomes with 1000 Genomes donors colored based on super population codes. EUR – European, EAS – East Asian, AMR – Admixed American, SAS – South Asian, AFR – African.

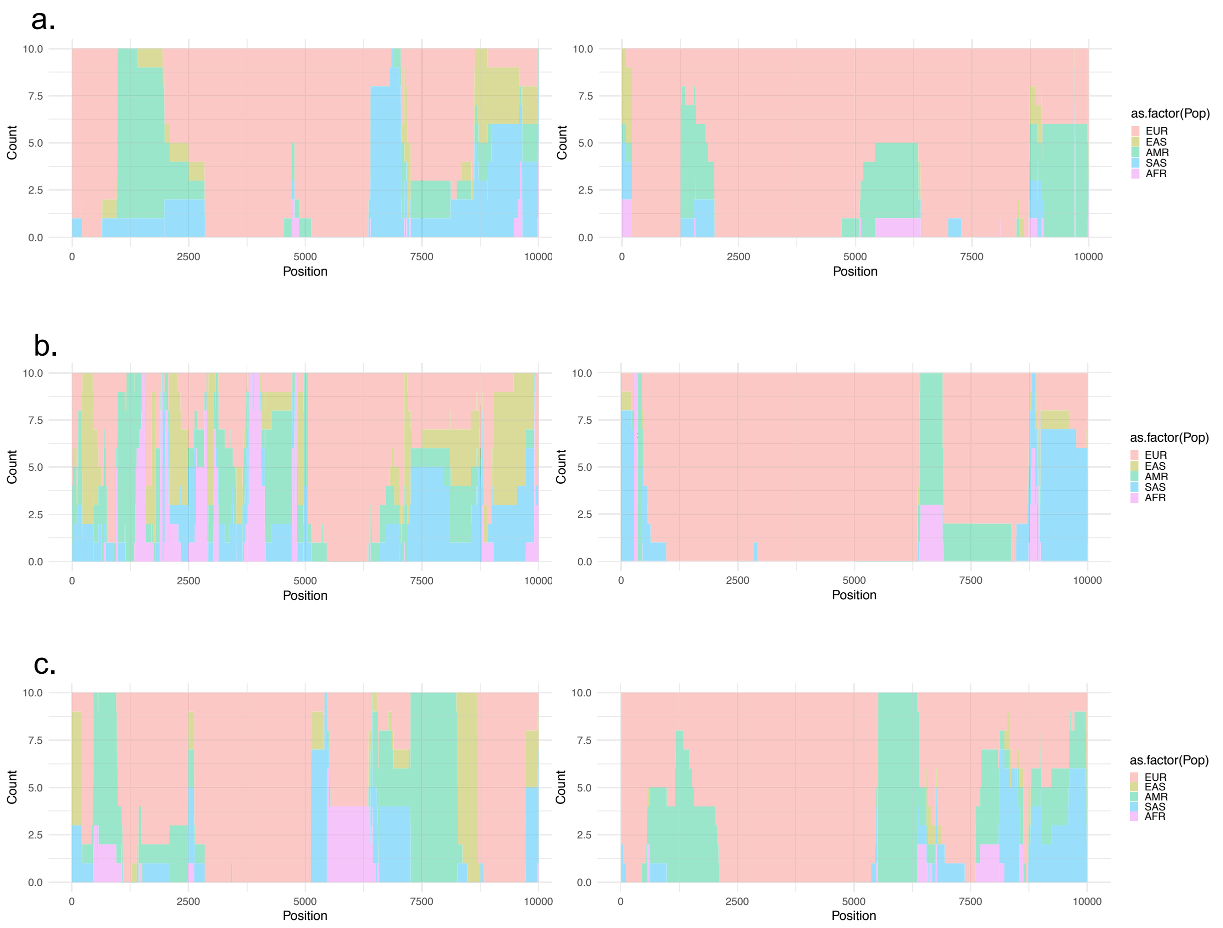

**Supplementary Figure 13. a)** Nearest neighbour adversarial accuracy ($AA_{TS}$) scores of artificial genomes generated from Estonian Biobank. Black line indicates the optimum value whereas values below the line indicate overfitting and values above the line indicate underfitting. **b)** Privacy loss. Test1 is a separate set of real Estonian genomes. Positive values indicate information leakage, hence overfitting.

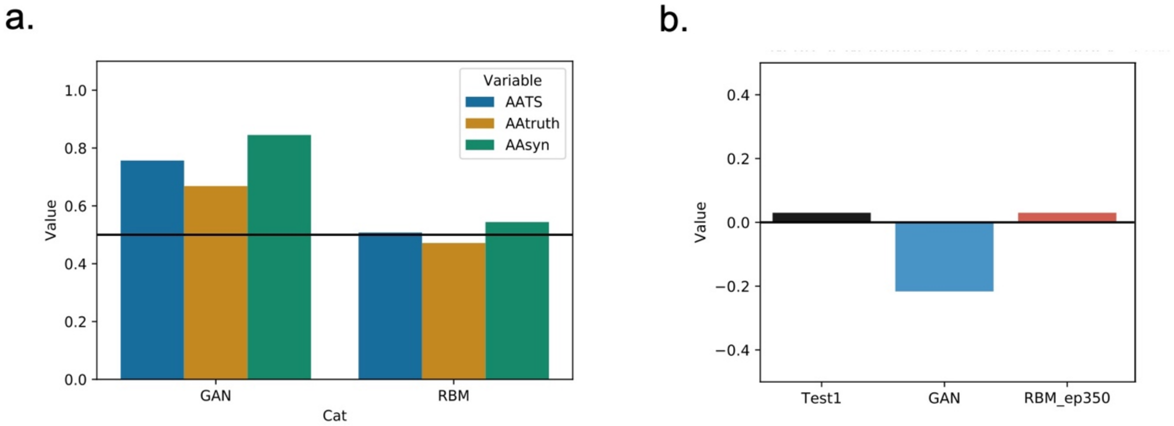

**Supplementary Figure 14.** Comparison of sites which are polymorphic in real genomes from Estonian Biobank but fixed in artificial genome counterparts generated via GAN and RBM models.

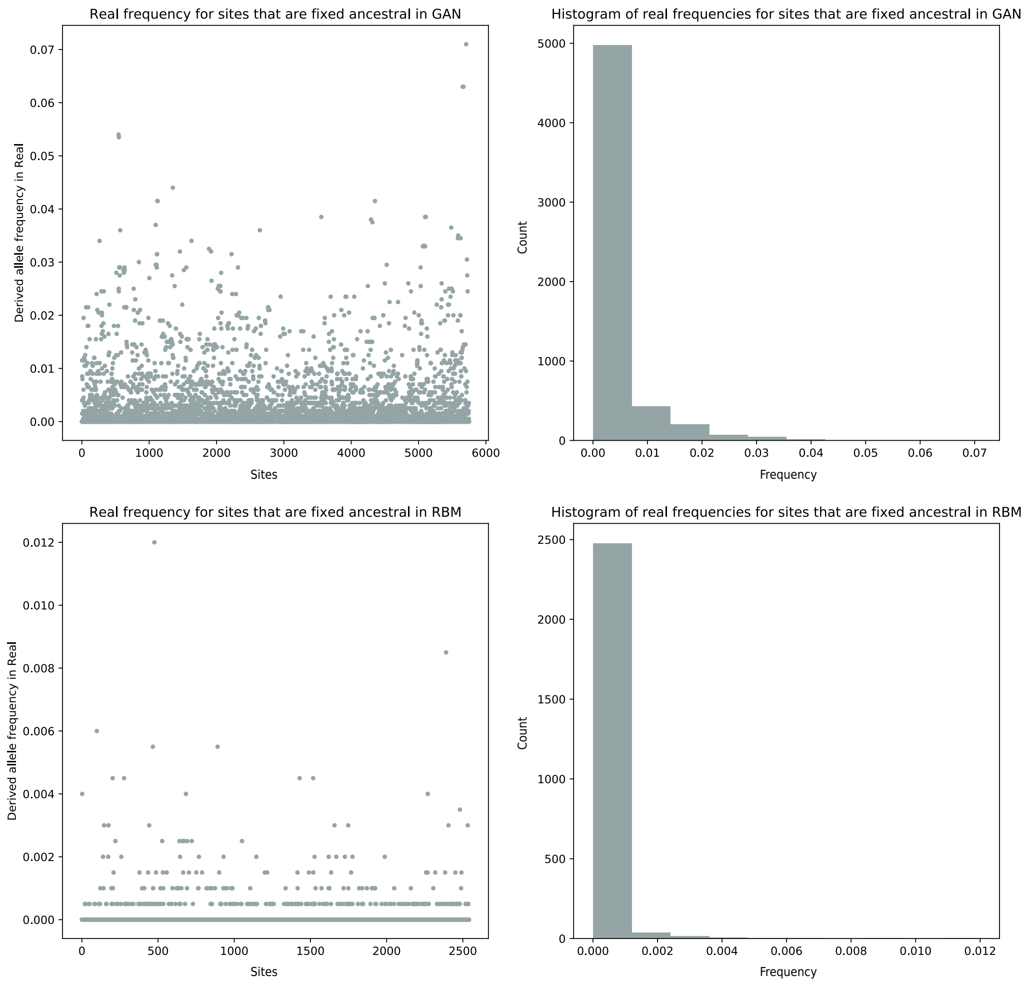

**Supplementary Figure 15.** Sensitivity tests for **a)** $AA_{TS}$ (scores over 0.5 indicate underfitting and below 0.5 indicate overfitting) and **b)** privacy scores (orange and red lines to mark the difference between RBM trained up to 350 and 690 epochs). All datasets consist of 2000 samples. Test1 and Test2 are real Estonian individuals who were not used in training. Mixed1 dataset has 1 real individual from the training dataset, Mixed2 has 10, Mixed3 has 50, Mixed4 has 100, Mixed5 has 500 and Mixed6 has 1000 individuals.

a.

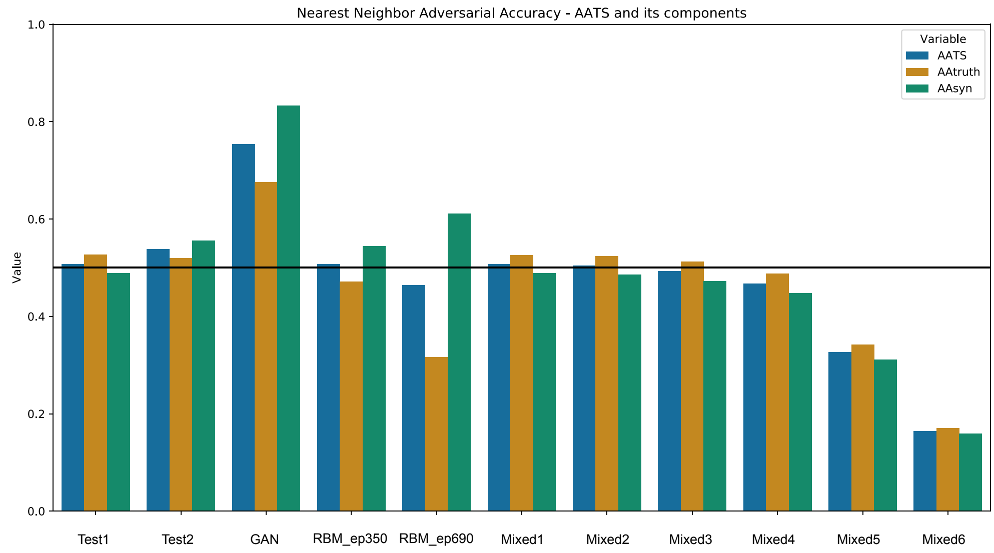


b.

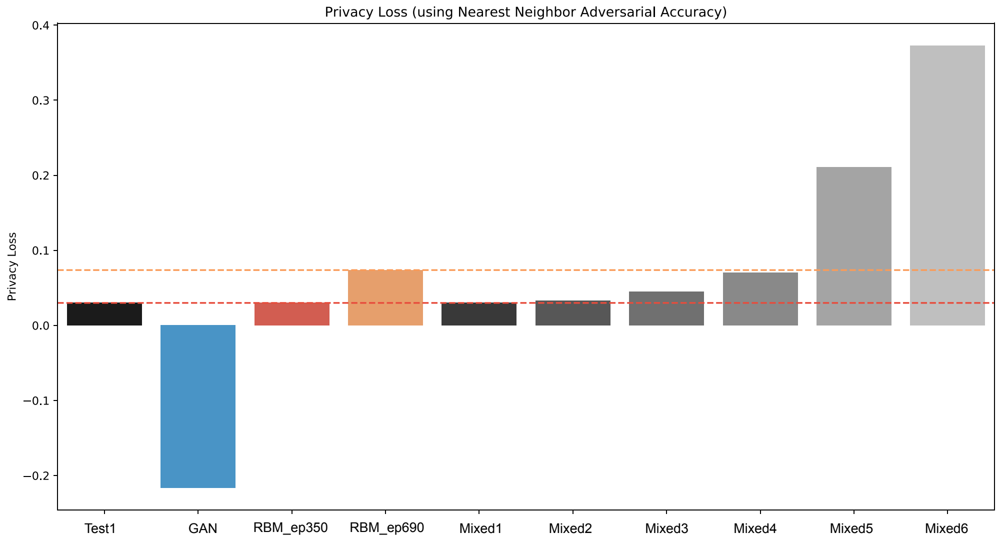

**Supplementary Figure 16.** Evaluation of $AA_{TS}$ scores of the GAN model for artificial Estonian genomes spanning **a)** 805 highly informative SNPs and **b)** dense 10K SNPs along with the total fixed sites for the outputs of epochs at 200 intervals.

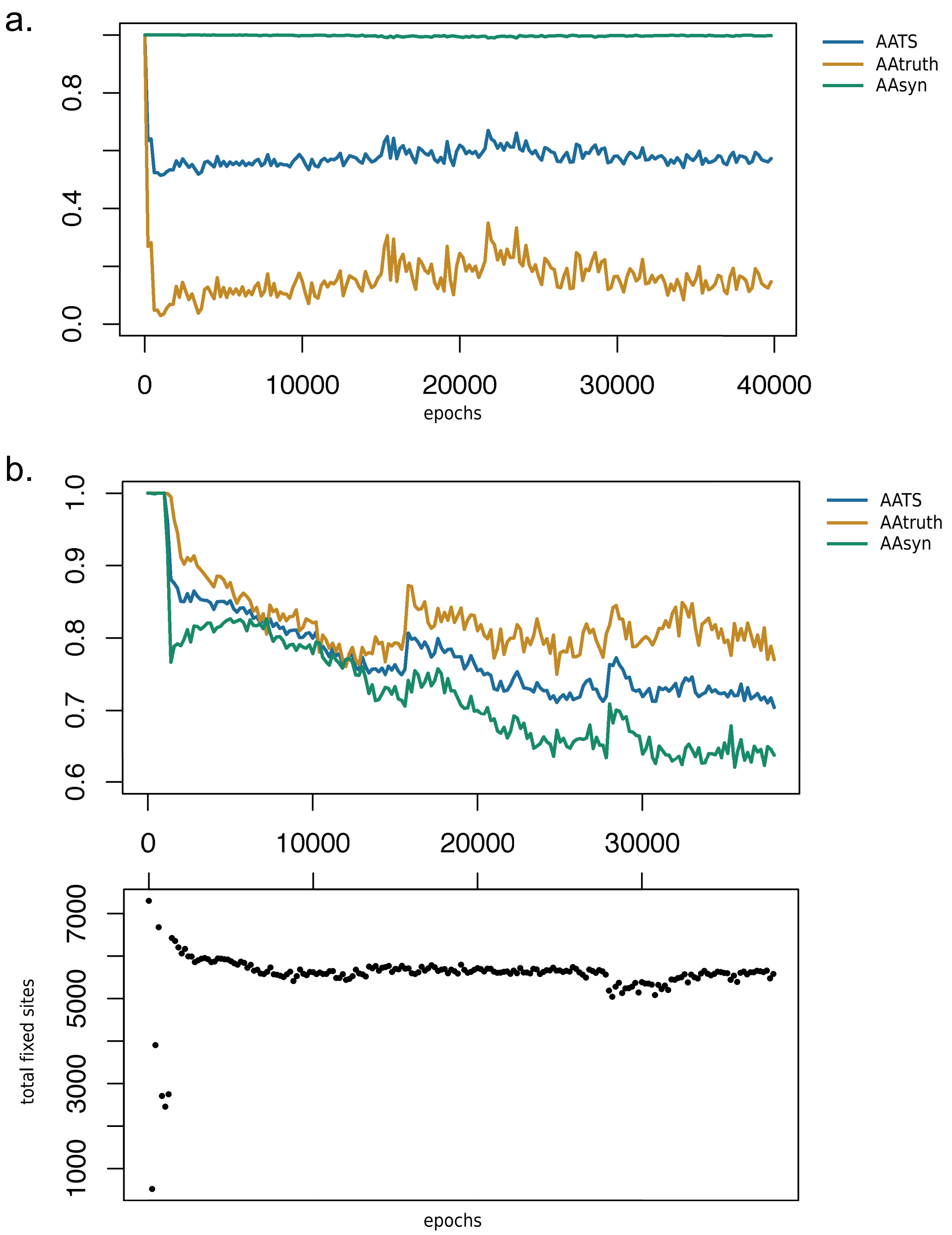

**Supplementary Figure 17.** Comparison of **a)** $AA_{TS}$ score and **b)** linkage disequilibrium of artificial genomes created via RBM model with sigmoid and ReLu activation functions.

a.

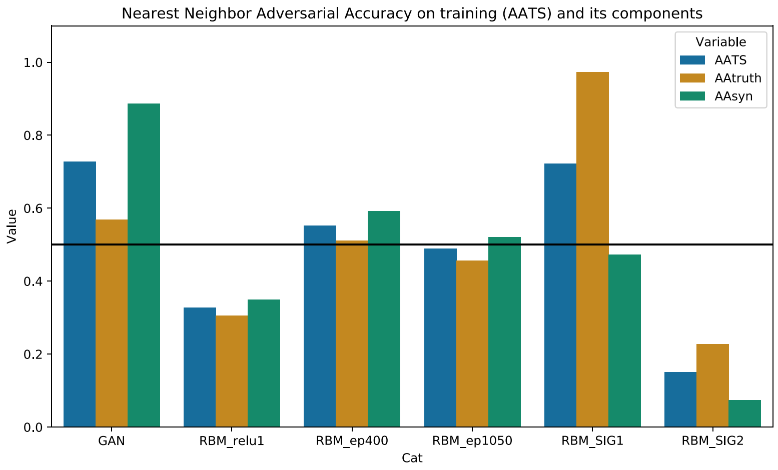


b.

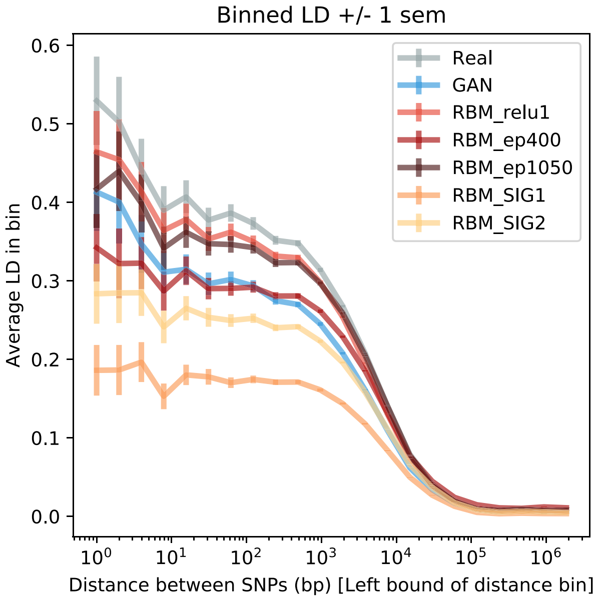
