## Supplementary material for "Creating Artificial Human Genomes Using Generative Models": Supplemetary Table

**Supplementary Table.** Genotype/phenotype contingency table for real and artificial Estonian genomes (AG). Ancestral allele "A" is associated with brown eye color and derived allele "G" is associated with blue eye color phenotype.

| **Real** | AA | AG | GG | Total |
| --- | --- | --- | --- | --- |
| Blue | 0 | 24 | 943 | 967 |
| Brown | 41 | 615 | 302 | 958 |
| Total | 41 | 639 | 1245 | 1925 |

| **AG** | AA | AG | GG | Total |
| --- | --- | --- | --- | --- |
| Blue | 9 | 95 | 778 | 882 |
| Brown | 28 | 377 | 638 | 1043 |
| Total | 37 | 472 | 1416 | 1925 |
