## Supplementary Text for "Creating Artificial Human Genomes Using Generative Models"

### Supplementary text: Restricted Boltzmann Machines

Yelmen, B et al.

September 2, 2019

As said in introduction the RBM consists of one visible layer and one hidden layer coupled by a weight matrix W. It is a probabilistic model of the joint distribution of visible $\{v_i, i = 1, \dots N_v\}$ and hidden variables $\{h_j, j = 1, \dots N_h\}$ of the form

$$P(v, h) = e^{-E(v,h)}$$

with

$$E(v, h) = \sum_{ij} W_{ij} v_i h_j + \text{bias terms}$$

Visible variables here are 0,1 as they represent ancestral or derived alleles, while the hidden variable type depends on the chosen activation function (sigmoid, RELU, ...). They are there to build dependencies among visible variables which by default are independent, via the interaction strength W. The weight matrix can be used in two different manners to interpret the learned model:

1. feature wise: for each hidden variable $j$ the vector $\{W_{ij}, i = 1, \dots N_v\}$ represents a certain combination of SNPs which, if activated, will contribute to activate or inhibits this feature $j$. These features are expected to be characteristic of the data structure (here the population structure) and the vector of feature activations should provide a suitable representation of individuals. If $N_v < N_h$ this corresponds to compressing the input representation.

2. direction wise: the SVD decomposition of W provides two sets of singular vectors with one corresponding to the visible space (visible axes) and the other one to the hidden representation (hidden axes). The one associated to the largest singular values offer the possibility to project the data in a low dimensional space. The dominant visible axes are expected to be similar to the principal component axes while dominant hidden axes are expected to produce more separable datapoints due to non-linear activation mechanisms. We used the latter (i.e. the projection into the hidden space) to perform our non-linear dimension reduction of the 1000Genomes data (see SM Figure).
